# Soft drug-resistant ovarian cancer cells invade via two distinct mechanisms utilizing myosin IIB

**DOI:** 10.1101/099655

**Authors:** Aastha Kapoor, Bhushan Thakur, Melissa Monteiro, Alakesh Das, Sejal Desai, Snehal Gaikwad, Amirali B. Bukhari, Pankaj Mogha, Abhijit Majumder, Abhijit De, Pritha Ray, Shamik Sen

## Abstract

The failure of chemotherapeutic drugs in treatment of various cancers is attributed to the acquisition of drug resistance. However, the invasion mechanisms of drug-resistant cancer cells remains incompletely understood. Here we address this question from a biophysical perspective by mapping the phenotypic alterations in ovarian cancer cells (OCCs) resistant to cisplatin and paclitaxel. We show that cisplatin-resistant (CisR), paclitaxel-resistant (PacR) and dual drug-resistant (i.e., resistant to both drugs) OCCs are softer and more contractile than drug-sensitive cells. Protease inhibition suppresses invasion of CisR cells but not of PacR and dual cells, suggesting protease-dependent mode of invasion in CisR cells and protease-independent mode in PacR and dual cells. Despite these differences, actomyosin contractility, mediated by the RhoA-ROCK2-Myosin IIB signaling pathway regulates both modes of invasion. Myosin IIB modulates matrix metalloproteinase-9 (MMP-9) secretion in CisR cells and nuclear squeezing in PacR and dual cells, thereby highlighting its importance as a potential therapeutic target for treatment of drug-resistant ovarian cancer cells.

**Financial Support:** Authors acknowledge financial support from IIT Bombay Healthcare Initiative, CSIR andDepartment of Biotechnology (Govt. of India) (Grant # BT/PR14658/MED/31/107/2010).AK and BTwere supported by fellowships from UGC and CSIR respectively (Govt. of India).

**Authors declare no competing financial interests**

Author contributions: AK, PR and SS designed the experiments. AK performed most of the experiments and analyzed the data.BT and SG developed drug resistant cell lines and did RT-PCR. MM performed western blot. AD performed AFM experiments. SD performed gelatin zymography. ABB, PM, AM and AD helped with live cell imaging experiments. AK and SS wrote the manuscript. All authors read and approved the final manuscript.

Summary statement: This study identifies drug-specific differences in the modes of invasion utilized by ovarian cancer cells, and demonstrates the role of myosin IIB in regulating both modes of invasion.

## Introduction

Epithelial ovarian cancer (EOC) is the leading cause of death in women suffering from gynaecological malignancies. Poor patient prognosis is associated with late detection and occurrence of drug resistance in ovarian cancer patients. Upon detection of EOC, cisplatin and paclitaxel represent the two widely used first-line therapy agents. However, post chemotherapy, nearly 90% patients develop resistance against these drugs. Drug resistance is associated with occurrence of epithelial to mesenchymal transition (EMT) across various types of cancers, imparting cells with enhanced aggressiveness and higher metastatic potential (Sarrió et al. 2008; Brabletz et al. 2005; Arumugam et al. 2009; Mitsumoto et al. 1998). Studies in ovarian cancer cells have shown that EMT leads to alterations in the biophysical properties of drug-resistant cells (Kim et al. 2013; Kajiyama et al. 2007). However, the functional implications of these changes vis-à-vis invasion remains to be understood. Further, given the distinct molecular mechanisms of action of cisplatin and paclitaxel, it is likely that drugs will induce distinct alterations. Therefore, understanding drug-specific changes which occur in these cells and their implications with regard to cancer invasiveness and mode of invasion represents an important step in refining our strategies for treating drug-resistant ovarian cancers.

For effective invasion through dense extracellular matrices (ECMs), cancer cells adhere to the matrix and make use of proteases to degrade the surrounding ECM thereby creating paths for their escape. This mode of invasion is referred to as the mesenchymal mode of invasion (Sabeh et al. 2009). Matrix metalloproteinases (MMPs) represent a family of 26 classes of proteases that are widely used by cancer cells for ECM degradation (Krakhmal et al. 2015; Tsai & Yang 2013). In drug-resistant ovarian cancers, upregulation of MMP-2 and MMP-9 is associated with poor patient prognosis(Fishman et al. 1997; Davidson et al. 1999; Alshenawy 2010; Schmalfeldt et al. 2001). In case of cisplatin-resistant head and neck carcinomas, upregulated MMP-7 and MMP-13 levels have been identified as biomarkers (Ansell et al. 2009). Though initial cisplatin treatment leads to inhibition of MMP-2 (Karam et al. 2010), development of resistance against anti-cancer drugs leads to upregulation of secreted MMPs. All these reports corroborate the occurrence of EMT in drug-resistant cells, suggesting that they use protease-dependent mesenchymal mode of invasion.

Upon inhibition of MMP activity, many cancer cells have been reported to transition to a rounded morphology. In this configuration, cells do not stop, but continue to migrate in a path finding mode wherein cells squeeze through existing pores in the matrix in an adhesion independent manner using actomyosin contractility (Wolf et al. 2013; Renkawitz et al. 2009; Fraley et al. 2010). This mode is referred to as the amoeboid mode of motility, and the transition from adhesion-dependent and protease-dependent mode of migration to adhesion-independent and protease-independent mode of migration is referred to as mesenchymal to amoeboidal transition (MAT). MAT is accompanied by change in cellular morphology (from spindle to round shape), abrogation of pericellular proteolysis and alterations in cytoskeletal organization(Friedl & Wolf 2003).

While the occurrence of EMT has been reported in drug-resistant ovarian cancers, it is unknown if drug-resistant cancers may undergo MAT. Also, the underlying molecular mechanism involved in invasion of drug-resistant cancer cells are unknown. In this study we address these questions by probing the biophysical properties of drug-resistant subtypes of OAW42, A2780 and TOV21G ovarian cancer cells, and correlating these properties with their mode of invasion. We show that though all drug-resistant subtypes (cisplatin-resistant, paclitaxel-resistant and dual drug-resistant) of OAW42 and A2780 cells, and TOV21G cells are more contractile and softer than drug-sensitive ovarian cancer cells. However, these cells differ in their proteolytic capabilities, with higher MMP activity of cisplatin-resistant cells which are similar to that of drug-sensitive cells, and negligible MMP activity in paclitaxel-resistant and dual drug-resistant cells suggesting that cisplatin-resistant, paclitaxel-resistant cells and dual drug-resistant use distinct modes of invasion, with cisplatin-resistant cells utilizing mesenchymal mode of invasion, and paclitaxel-resistant and dual drug-resistant cells utilizing amoeboidal mode of invasion. Interestingly, in spite of the different invasion strategies utilized by these cells, cellular contractility is modulated in all drug-resistant subtypes via the RhoA-ROCK2 signaling pathway by regulating non-muscle myosin IIB activity. In TOV21G cells which utilize mesenchymal mode of invasion, myosin IIB knockdown inhibits invasiveness by inhibiting MMP secretion. In contrast, in dual drug-resistant OAW42 cells which utilize amoeboidal mode of invasion, nuclear deformability is compromised leading to less invasion. Taken together, our results suggest that the same molecular players are able to direct differential modes of invasion in cells exhibiting resistance to two different cytotoxic drugs and therefore could be potential therapeutic targets.

## Materials and Methods

### Cell Culture

A2780, OAW42 and TOV21G human epithelial ovarian cancer cell lines were obtained from ACTREC (Advanced Centre for Treatment, Research and Education in Cancer, Mumbai, India) and drug-resistant derivatives of A2780 and OAW42 cells were developed by exposing them to cyclic drug treatment. Cisplatin (CisR), paclitaxel (PacR) and dual-drug (Dual) resistant models were developed by subjecting A2780/OAW42 (1 × 10^6^cells) to incremental doses of cisplatin or/and paclitaxel (Sigma-Aldrich, cisplatin-P4394, paclitaxel-T7191) for a period of 6 months with each dose repeated for three cycles. Based upon the drug-type used, cells were categorized as CisR (cisplatin-resistant), PacR (paclitaxel-resistant) and dual (resistant to both cisplatin and paclitaxel). TOV21G cells were not subjected to any drug treatment since they are intrinsically cisplatin-resistant in nature (Thakur & Ray 2016). CisR, PacR and Dual cell lines were maintained using IC_50_ concentrations of cisplatin or/and paclitaxel drugs (Singh et al. 2014). Cells were grown in Dulbecco’s Modified Eagle Medium (Gibco,cat#12800-017) supplemented with 10% fetal calf serum, penicillin/streptomycin incubated at 37°C with 5% CO2. For drug related studies, the broad spectrum MMP inhibiting drug GM6001 (Calbiochem, cat#364206, 5-10μM) and myosin II inhibiting drug blebbistatin (Abcam,cat#120491, 10-20μM) were added 4 hours after seeding the cells to allow cell substrate attachment, except in case of collagen degradation assay, in which they were added half an hour post seeding. All cell lines A2780, OAW42 and TOV21G were authenticated by STR profile analysis. Genomic DNA was supplied to DNA Labs India (govt. certified lab for forensic studies) andDNA test report shows >80% profile match.

### Extracellular Matrix (ECM) Degradation

For assessing ECM degradation capability of cells, the collagen degradation assay was performed wherein drug-sensitive and drug-resistant ovarian cancer cells were plated on fluorescently labelled collagen I coated glass coverslips. Glass coverslips were incubated with rat tail collagen I (Sigma, cat#C3867)at a concentration of 10 μg/cm^2^ and incubated overnight at 4°C to allow passive adsorption. To prevent non-specific binding, the coverslips were subsequently blocked with 1% Pluronic F127 (Sigma, cat#P2443) for 10 minutes and rinsed with PBS. Collagen was stained with mouse polyclonal primary antibodies (Merk, custom made) at 1:500 overnight and counterstained with anti-mouse secondary antibody (Bangalore Genei) at 1:1000 for 2 hours. Collagen degradation was assessed after culturing cells for 6 hours to allow for visible degradation. Quantification of the degraded area was performed using ImageJ (NIH).

### Immunofluorescence (IF)

Drug-sensitive and drug-resistant ovarian cancer cells were fixed with 4% paraformaldehyde (Sigma,cat#158127) in PBS for 20 mins, followed by blocking with 1% bovine serum albumin (BSA) for 1 hour at room temperature. Cells were incubated with primary antibodies at 4°Covernight followed by incubation with secondary at room temperature (RT) for 2hours. Varying dilutions of primary antibodies were used in case of vinculin (1:400, Abcam, cat#18058), pMLC (1:200, CST, cat#3674S), NMM IIA (Abcam, cat#55456), NMM IIB (1:500, Abcam, cat#684).Secondary antibodies (Bangalore genei) were added at a dilution of 1:1000, unless otherwise specified. Phalloidin (Invitrogen) and DAPI (Sigma) dyes were used at 1:500 for staining actin filaments and nucleus, respectively. Samples were mounted with Eukit (Sigma, cat#03989) and imaged with confocal microscope (OlympusIX81). Image processing and protein quantification was done using Image J (NIH) as described elsewhere(Kapoor & Sen 2012). Briefly, to quantify the amount of vinculin localization at focal adhesions (FA), images were thresholded, and thresholded vinculin area for each cell was divided by the total cell area (using the F-actin signal) to calculate the percentage of FA area per cell. In order to quantify pMLC levels, mean signal intensity of every cell was measured. At least 30 cells across three independent experiments were analyzed for vinculin and pMLC protein quantification, respectively.

### Spreading, de-adhesion assay and nuclear height measurement

For spread area analysis, drug-sensitive and drug-resistant cells were plated on 0.1 μg/cm^2^ collagen-coated coverslips and imaged after 24hrs incubation using inverted microscope (Olympus IX71). Quantification was done by manually tracing the cell perimeter to measure enclosed area using Image J (100 cells per condition across three independent experiments). For trypsin deadhesion experiments, time-lapse videos of trypsin treated cells was acquired (Sen & Kumar 2009). Briefly, cells were washed with PBS, incubated with 1X warm trypsin (Himedia) and time-lapse video was taken at an interval of 1 sec until the cells rounded up. As above, atleast 50 cells across three independent experiments were analyzed per condition. For measuring nuclear height of cells, live cells were stained with DAPI at 1:1000 concentration for 3mins followed by z-stack imaging with multiphoton confocal microscope (Zeiss LSM780). Thorough PBS washes (thrice for 2 mins each) were given prior to imaging to remove excess DAPI stain. Z-stack imaging was done spanning 60μm of vertical height region followed by orthogonal reconstruction and nuclear height assessment using Zen Blue software.

### Atomic Force Microscopy (AFM)

Force curves were obtained with an Asylum MFP3D AFM (Asylum Research, CA) mounted on a Zeiss epifluorescence microscope and indented using a pyramid-tipped probe (Olympus) with nominal spring constant of 20 pN/nm(MacKay & Kumar 2012). Actual spring constants were determined using thermal calibration method. Cells were cultured overnight on collagen-coated 60mm plates and indented. Cortical stiffness was determined by fitting each force curve with a modified Hertzian model of a pyramid indenting a semi-infinite elastic solid(MacKay & Kumar 2012). At least 100 cells were indented per condition across at least three independent experiments. Matlab software was used to calculate young’s modulus (*E_c_*) by running a customized code.

### Western Blotting

Whole cell lysates from drug-sensitive and drug-resistant ovarian cancer cells were lysed using RIPA buffer (Sigma, cat#R0278) containing 1× protease inhibitor cocktail and phosphatase inhibitor cocktail 3 (Sigma). The lysed cells were centrifuged at 10,000 ×g for 20 mins, the supernatant was collected and the protein concentration was quantified using Bradford assay (Himedia, cat#ML106). The protein samples were boiled at 95°C for 5 min in 1×Laemmli buffer with β-mercaptoethanol. Protein samples were loaded onto a 15% SDS-polyacrylamide gel and transferred onto nitrocellulose membranes (BioTrace NT Membrane-Pall). This was followed by blocking in 5% BSA in TBST buffer (10 mMTris–HCl, pH 8.0 containing 150 mMNaCl and 0.1% Tween20). Membranes were washed three times in TBST for 15 mins each and incubated overnight at 4°C with specific antibodies. Antibodies diluted in TBST buffer at 1:400 for anti-vinculin antibody (Abcam, cat#18058), 1:1000 for phospho-myosin light chain 2 antibody (CST, cat#3674S), 1:500 for non-muscle myosin IIA/B antibody and 1:2500 for anti-GAPDH antibody were used as primary antibodies. Membranes were washed five times with TBST buffer and incubated with horseradish peroxidase-conjugated secondary antibodies for 1h at room temperature. Membranes were washed three times in TBST and developed using a chemiluminescence detection system (Pierce ECL Western Blotting Substrate-ThermoScientific).

### Gelatin zymography

Drug-sensitive and drug-resistant ovarian cancer cells (5×10^4^ cells) were grown in 750μl of media in 24 well plates. After 36 hrs of incubation, conditioned media was collected and centrifuged at 10,000 rpm for 10 min to remove cells (if any) and subjected to gelatin zymography. In brief, media volumes containing equal amounts of protein were mixed with 6X Laemmli non-reducing buffer, incubated for 15 min at room temperature. Samples were then electrophoresed on 8% SDS-PAGE gel containing 1 mg/ml gelatin. After electrophoresis, the gels were washed twice with renaturation buffer (2.5% triton X-100) for 30 min each, followed by rinsing 3 times with reaction buffer (50 mMTris–HCl buffer (pH 7.6), 5 mM CaCl2, 200 mMNaCl) and incubated in fresh reaction buffer overnight at 37 °C. The gels were stained with 0.5% Coomassie brilliant blue G-250 solution for 30 min and destained with 7.5% acetic acid solution containing 10% methanol. Areas of gelatinase activity were detected as clear bands against the blue-stained gelatin background, which was quantified by densitometric analysis using ImageJ software.

### Preparation of 3D collagen sandwich

3D collagen sandwich was prepared on functionalized cover slips(Artym & Matsumoto 2010). Coverslips were treated with 0.1N NaOH for 15mins, followed by silane treatment for 15mins, followed by final treatment with 0.5% glutaraldehyde for 15 mins, to functionalize them. Using rat tail collagen I, 3D collagen cocktail was prepared according to manufacturer’s guidelines (Corning, cat#354236). Briefly, working volume of 3D collagen was calculated for final concentration of 1 mg/ml. The solution was neutralized with equal volumes of 100 mM of HEPES buffer. Next, 7.5% NaHCO_3_ and 0.1N NaOH was added to the neutralised solution. Cocktail was then mixed homogeneously and the entire set up kept on ice to prevent rapid gelation. 30μl of final mix was layered onto each functionalized cover slip and incubated for 30 mins at 37°C. Ovarian cancer cells were plated on this pre-formed layer and allowed to settle for 2 hours. Subsequently, a second layer of 3D collagen was added on top of the adhered cells at two concentrations 0.5 mg/ml and 1 mg/ml, and the samples put at 37°C to allow the top layer to polymerize. Media was added after the top layer polymerized. For viewing cell entrapment within the sandwich gel, gels were fixed and stained as described elsewhere (Artym & Matsumoto 2010). Briefly, 4% PFA (warmed at 37°C) was used to fix cells for 20 mins, followed by permeablization with 0.5% Triton X-100 for 10mins. Blocking was done using 1%BSA for 30 mins, followed by staining collagen with primary antibody at 1:1000, for 2 hrs, at RT and secondary antibody at 1:1500 for 2 hrs, at RT. Nuclei staining was done using DAPI at 1:1000 for 3 mins. Each step was followed by three PBS washes of 15mins each. Imaging was done using multiphoton confocal microscope (Zeiss LSM780) and orthogonal reconstruction was done using Zen Black software. Set up was used for tracking cell invasion, time lapse images were acquired every 20 mins for a duration of 15 hours. As before, cell speed was measured using the manual tracking plug-in of ImageJ. Atleast 60 cells were analyzed per condition and the experiment was repeated thrice.

### Semi-quantitative RT-PCR analysis

Reverse transcriptase coupled PCR was performed following manufacturers protocol. Briefly, total RNA was extracted from cultured cells using RNease kit (Qiagen,cat#74104). One microgram of total RNA was reverse transcribed using cDNA synthesis kit (Invitrogen,cat#11904-018). GAPDH was used as an internal control.

**Table 1.**
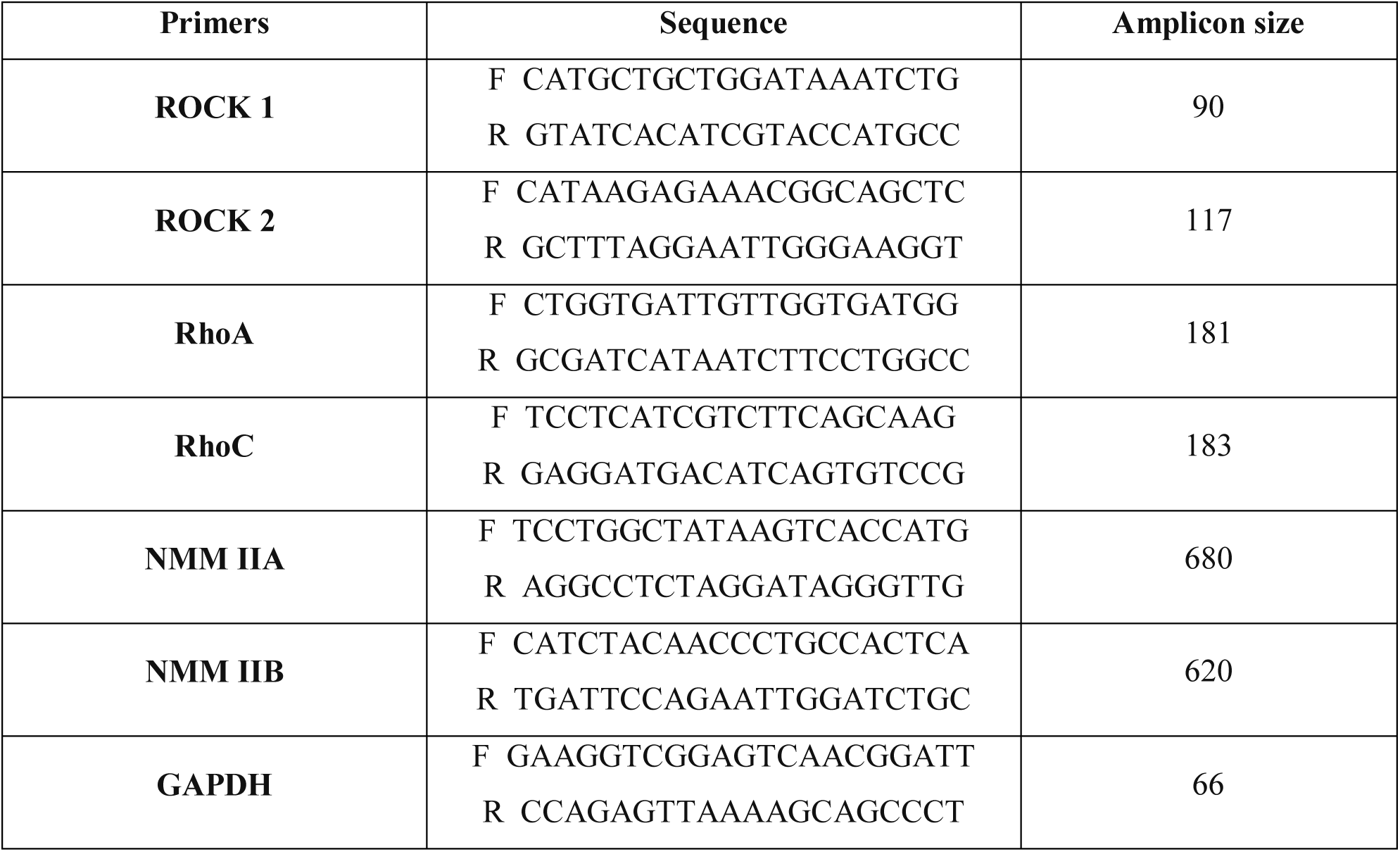
Contains the pair of primer sequences for different genes along with their amplicon size used during RT-PCRs.

### shRNA based non-muscle myosin IIB (NMM IIB) knockdown

For NMM IIB knockdown studies, 10^4^ cells were seeded in 96 well plates (in triplicates) and infected with lentivirus encoding NMM IIB shRNA (Sigma, target sequence: CAACATTGAAACATACCTTCTCGAGAAGGTATGTTTCAATGTTG) and empty vector (Sigma,cat#SHC001V)at a multiplicity of infection (MOI) of 3 in culture medium supplemented with hexadimethrine bromide (8 μg/ml; Sigma). The following day, the medium of the infected cells was aspirated and replaced with fresh medium for stabilization. Selection for transfected cells was started on third day by adding media containing puromycin (1μg/ml). Cells were supplied with fresh puromycin media every second or third day from the commencement to enrich resistant cells. After twelve days of selection, resistant cells were collected and proliferated to obtain a stably transfected population. NMM IIB knockdown was confirmed by western blotting.

### Statistical Analysis

Unpaired student’s t-test was used to find the statistical significance of the data. P values below 0.05 were considered to be significant for all experiments, unless otherwise specified. Origin 8.0 software was used to perform statistical analysis and for performing curve fitting wherever required.

## Results

### Drug-resistant cells are more contractile than drug-sensitive cells

Drug-resistant ovarian cancer cells have been reported to undergo EMT, and possess high metastatic potential (Kim et al. 2013; Kajiyama et al. 2007). However, their mechanism of invasion is not fully clear. As a first step towards investigating the molecular mechanisms of invasion in drug-resistant ovarian cancer cells, we performed morphometric analysis of drug-sensitive A2780 and OAW42 serous carcinoma cells, and their drug-resistant counterparts developed by exposing them to either cisplatin (CisR), paclitaxel (PacR) or a combination of both (dual). Experiments were also performed with TOV21G endometroid carcinoma cells which are naturally resistant to cisplatin. Even though both OAW42 and A2780 are serous grade carcinoma cells, they differ inherently in their cell sizes, with OAW42 being bigger (12-62 μm^2^) and A2780 being smaller (2-6 μm^2^) in spread area (**Fig. S1A,B**). TOV21G cells were intermediate in size (16-20 μm^2^). Moreover, while cisplatin and paclitaxel treatment induced cell rounding in both OAW42 and A2780 cells, cells treated with both these drugs together exhibited morphology similar to that of drug-sensitive cells.

To probe if cell rounding induced by cisplatin and paclitaxel treatment was due to alterations in the number of focal adhesions and/or levels of focal adhesion proteins, cells were probed for expression levels and localization of the focal adhesion protein vinculin. Compared to drug-sensitive cells, vinculin expression was lower in all the drug-resistant cell types (**Fig. 1B**). However, vinculin localization at focal adhesions was found to be sensitive to the type of drug treatment, with prominent focal adhesions detected in both drug-sensitive, cisplatin-resistant (CisR) cells and TOV21G cells, but absent in paclitaxel-resistant (PacR) cells and dual drug-resistant (dual) cells (**Fig. 1A**).

**Figure 1:**
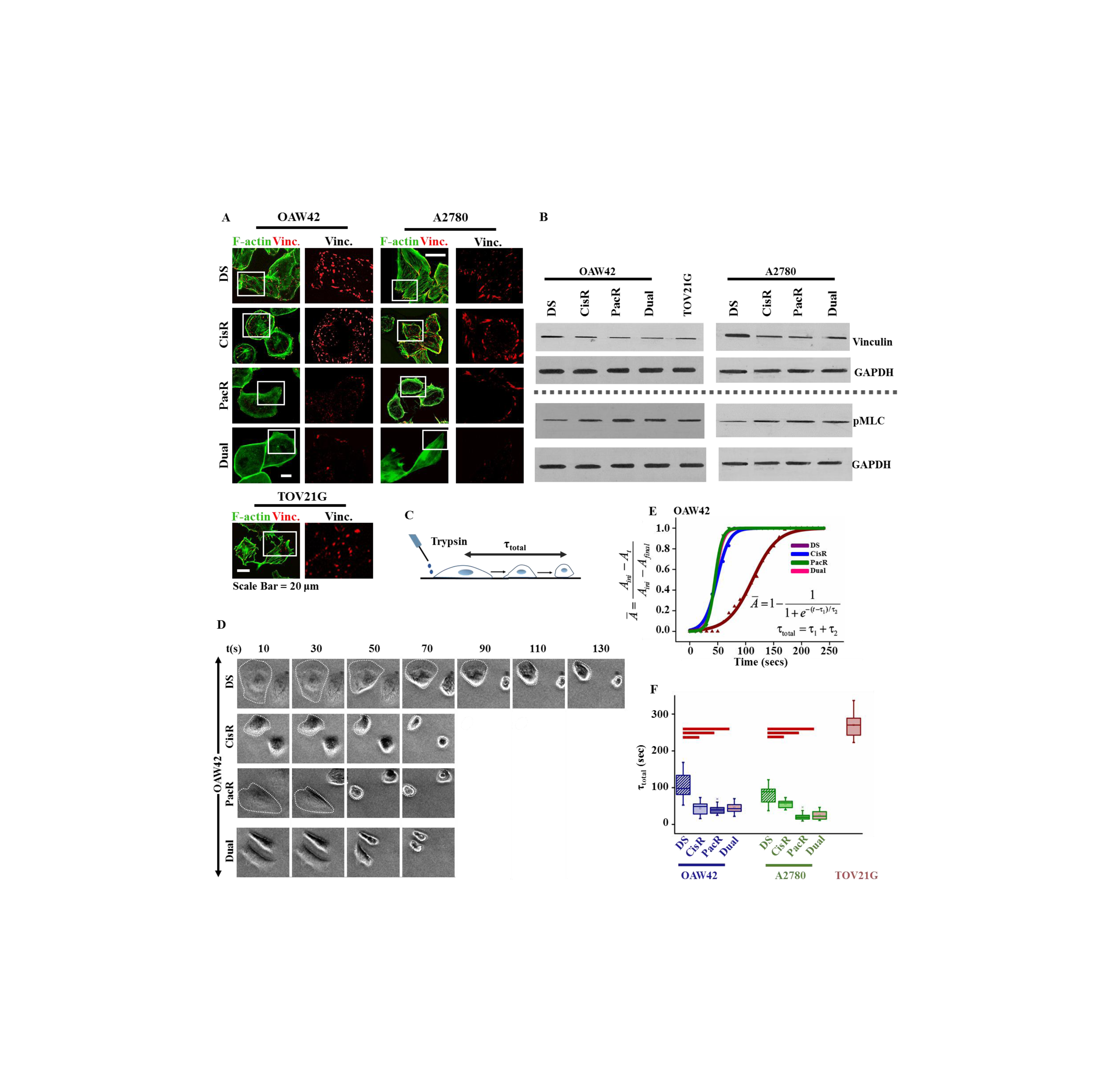
Drug-resistant ovarian cancer cells are more contractile compared to drug-sensitive cells. **(A)** Representative immunofluorescence images of focal adhesion (FA) localized vinculin (Scale bar = 20μm) in drug-sensitive (DS) and drug-resistant OAW42 and A2780 cells, and TOV21G cells. **(B)** Western blots of total vinculin levels and phosphorylated myosin light chain (pMLC) levels in DS, cisplatin-resistant (CisR), paclitaxel-resistant (PacR) and dual drug-resistant (dual) cells. GAPDH was used as a loading control. **(C)** Schematic of trypsin de-adhesion assay for assaying cell contractility. In this assay, adherent cells are incubated with warm trypsin and their rounding kinetics tracked till they round up but remain attached to their substrates. τ_total_ represents the total rounding time.**(D)**Representative phase contrast, time-lapse images of de-adhesion kinetics of DS, CisR, PacR and dual cells. PacR and dual cells were found to exhibit fastest de-adhesion dynamics (Scale bar = 45μm).Refer to supplementary video1.**(E)** De-adhesion dynamics was quantified by tracking the fraction change in area 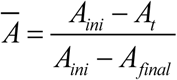 as a function of time. In this expression, *A_ini_* and *A_fin_* represent the initial and final cell area for a given cell, and *A_t_* represents the cell area at time *t*. The de-adhesion curves are fitted with the expression 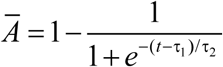 to obtain the de-adhesion time constants τ_1_ and τ_2_, respectively. The total de-adhesion time (τ_total_) is the sum of these two time constants. **(F)** Plot of average total deadhesion time of DS, CisR, PacR and dual cells. At least 50 cells across three independent experiments were analyzed for each condition. Error bar indicates SEM. Red bars indicate statistical significance (*p < 0.05).

While cisplatin and paclitaxel perturbed focal adhesion organization differentially, robust increase in phosphorylated myosin light chain (pMLC) was observed across all the drug treatments, indicative of increased actomyosin contractility in drug-resistant cells (**Fig. 1B**). To test this functionally, trypsin de-adhesion assay was performed wherein adherent cells are incubated with warm trypsin and their rounding kinetics tracked until cells are rounded but remain attached to their substrates (**Fig. 1C inset**). Using a variety of different cell types, we had previously demonstrated that de-adhesion timescales (τ_1_ and τ_2_) are directly influenced by actomyosin contractility with higher cellular contractility leading to faster de-adhesion(Sen & Kumar 2009; Kapoor & Sen 2012; Sen et al. 2009)(**Fig. 1D**). Consistent with higher pMLC levels in drug-resistant cells, the total de-adhesion time τ_total_ (=τ_1_ + τ_2_) of drug-resistant cells(PacR, CisR and dual) was lower (i.e., faster de-adhesion) compared to DS cells (**Fig. 1 C-E, Supp. Video. 1**). In comparison to OAW42 and A2780 cells, TOV21G exhibited delayed de-adhesion. Collectively, our results suggest that drug-resistant cells possess higher contractility compared to drug-sensitive cells.

### Drug-resistant cells are softer than drug-sensitive cells

While cell softness has been associated with increased metastatic potential in ovarian cancer cells (McGrail et al. 2014), recent reports have reported that drug-resistant ovarian cancer cells are stiffer than drug-sensitive cells (Sharma et al. 2014). To evaluate how stiffness of drug-resistant cells compared with those of drug-sensitive cells, we performed AFM stiffness measurements on single cells. The first 500 nm of the force-indentation curves were fit using Hertzian model to estimate cortical stiffness (*E*) of drug-sensitive and drug-resistant cells (MacKay & Kumar 2012)(**Fig. 2A**). Surprisingly, in contrast to recently published reports, CisR, PacR and dual cells were all found to be significantly softer than DS cells (**Fig. 2B**). TOV21G cell stiffness was also found to be comparable to that of the other drug-resistant cells. Collectively our results suggest softening of ovarian cancer cells upon acquisition of drug-resistance.

**Figure 2:**
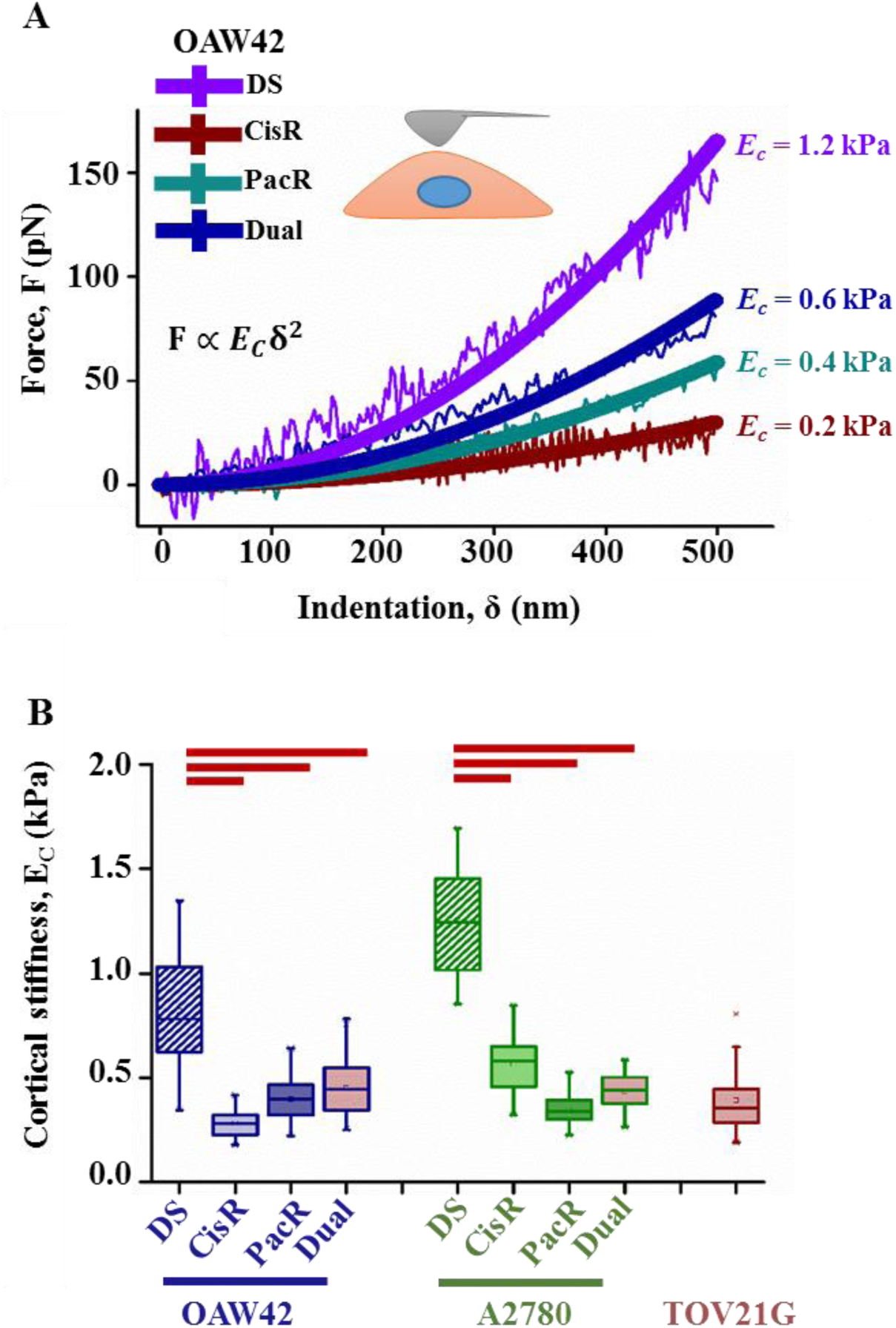
Drug-resistant cells are softer than drug-sensitive cells. **(A)**Probing of cortical stiffness using Atomic Force Microscope (AFM). After 24 hours in culture, cells were probed with a soft pyramidal tip for measuring their cortical stiffness. Representative force-indentation curves obtained by probing DS, CisR, PacR and dual OAW42 cells. Experimental curves were fit using the Hertz theory to estimate cortical stiffness (*E_c_*) of cells.**(B)** Cortical stiffness of DS, CisR, PacR and dualsubtypes of OAW42 and A2780 cells, and TOV21G cells. Data represented as mean ± SEM (n = 100 cells for each condition, experiment repeated thrice). Red bars indicate statistical significance (*p < 0.05).

### Cisplatin-resistant and paclitaxel-resistant cells use distinct modes of invasion, both modulated by actomyosin contractility

To further probe if drug-resistant cells utilized similar or different modes of invasion, ECM remodeling experiments were performed using collagen degradation assay, wherein cells were cultured on pre-labelled collagen-coated substrates (10 μg/cm^2^) for a period of 6 hours and degradation was assessed from the presence of dark areas (i.e., degraded spots) underneath the cells. While similar degradation profiles were observed in DS and CisR cells, negligible degradation was observed in PacR and dual cells (**Fig. 3A**). It was not possible to perform degradation experiments with TOV21G cells as they stained with collagen-specific antibodies, interfering with capture and analysis of degraded collagen zones, i.e., the degraded zones were masked by the signal from the TOV21G cells (**Fig. S2**). The prominent degraded areas underneath the cells in DS and CisR cells could be attributed to higher MMP-9 activity, as revealed by gelatin zymography experiments (**Fig. 3B**). MMP-9 activity was also detected in TOV21G cells. In contrast, very low MMP activity was detected in PacR and dual cells, consistent with the absence of degraded areas.

**Figure 3:**
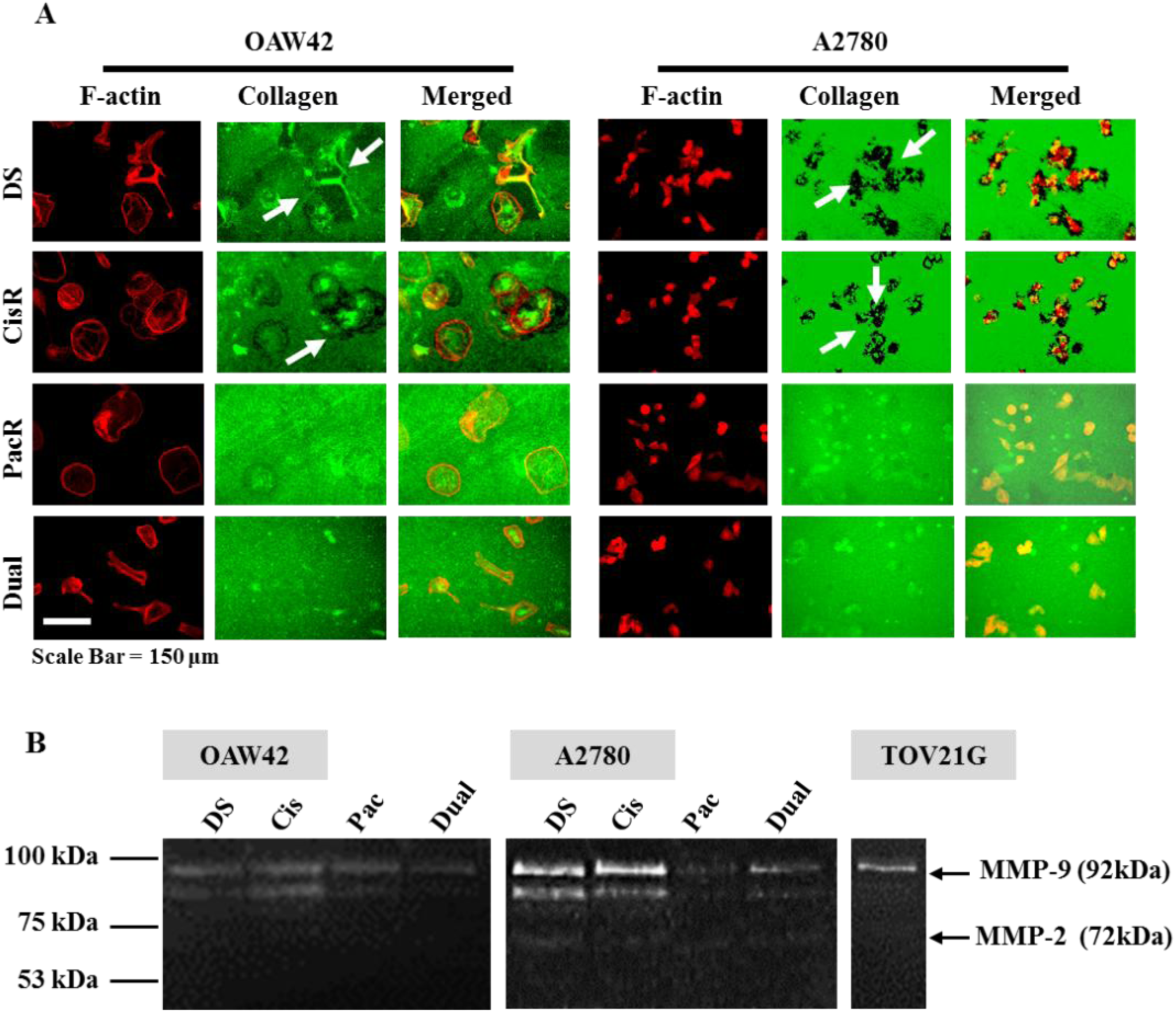
ECM degradation ability and MMP activity ofdrug-sensitive and drug-resistant cells. **(A)** Assessment of ECM degradation capabilities of cells using collagen degradation assay. In this assay, glass coverslips are coated with collagen I at a concentration of 10 μg/cm^2^. Subsequently, the coverslips are washed to dislodge loosely attached collagen, and then labeled with fluorescent antibody. Cells were cultured on these substrates for duration of 6 hours. Subsequently, cells were fixed and stained with phalloidin to visualize the cell boundaries. Collagen degradation was assessed by quantifying the degraded areas (i.e., areas devoid of fluorescent signal) underneath the cells (Scale bar = 100μm).**(B)** Quantification of proteolytic activity of DS, CisR, PacR and dual cells. Gelatin Zymography was performed using conditioned media from DS, CisR, PacR and dual cells after 21 hours in culture. Prominent MMP-9 bands were detected in DS and CisR cells but not in PacR and dual cells.

The combination of ECM degradation and zymography experiments are indicative of protease-dependent mode of migration utilized by DS and CisR cells, and protease-independent migration by PacR and dual cells. To probe this in a more *in vivo*-mimetic scenario, we performed invasion assay using the sandwich gel setup where cells are confined within two layers of collagen gels (**Fig. 4A**). In this setup, cells remain at the interface (**Fig. 4B**) and migrate laterally, with the top collagen gel providing steric hindrance. A 1 mg/ml collagen gel was chosen as pore size of these gels (~7 μm^2^) (**Fig. S3A**) was found to be lesser than nuclear height, and hence expected to provide hindrance for lateral cell migration. Experiments were performed in the absence and presence of the broad spectrum MMP inhibitor GM6001, and the contractility inhibiting drug blebbistatin. Though the lateral invasion speeds were comparable across the different cell types, GM6001 perturbed cell speed to different extents. In DS, CisR and TOV21G cells, where robust MMP activity was detected, GM6001 treatment significantly inhibited lateral invasion (**Fig. 4C, Supp. Videos. 2-4**). In contrast, lateral invasion was minimally affected in PacR and dual cells, where collagen degradation was minimal. Blebbistatin treatment, which inhibits cell contractility, nearly abolished lateral invasion across all the cell types, in both drug-sensitive and drug-resistant cells. Together, these findings indicate that cisplatin and paclitaxel-resistant cells utilize distinct modes of invasion, and are indicative of a central role for contractility in regulating both modes of invasion.

**Figure 4:**
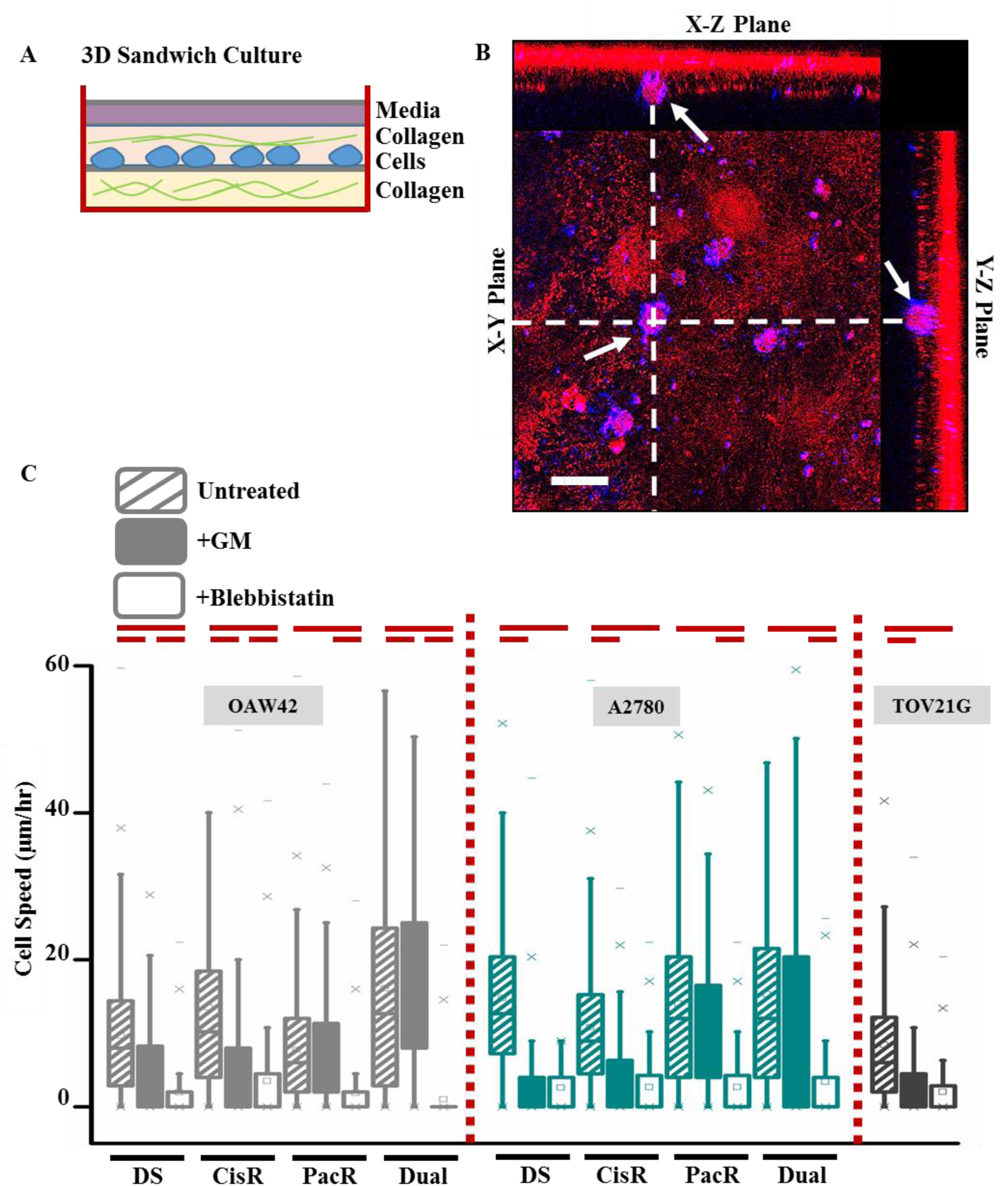
Cisplatin-resistant and paclitaxel-resistantcells utilize distinct modes of invasion in a contractility-dependent manner. **(A)** Schematic of 3D sandwich invasion assay setup used for quantifying cell invasiveness. In this assay, cells were grown between two layers of 3D collagen I at a concentration of 1mg/ml. Cells were tracked every 20 mins for a duration of 15 hours. For drug studies, media was supplemented with GM6001 and blebbistatin, respectively.**(B)** Orthogonal view of sandwich gel setup acquired with confocal microscope. Collagen was labelled with red fluorophore tagged antibody and cells were stained with DAPI for visualizing cell nuclei(Scale bar=10 μm). **(C)** Quantitative analysis of cell speeds of DS, CisR, PacR and dual cells in the presence and absence of drugs. (n≥30cellsper condition across three independent experiments), Red bars indicate statistical significance (*p< 0.05) assessed using Student’s t test. (Refer to supplementary videos 2-4).

### Myosin IIB is essential for both protease-dependent and protease-independent modes of invasion

To identify the molecular players regulating cellular contractility in drug-resistant cells, transcript profiling of drug-sensitive and drug-resistant cells was performed with a particular focus on molecules pertaining to the Rho signaling pathway, which has been implicated in regulating actomyosin contractility and promoting cell movement and metastasis (Karlsson et al. 2009). Between drug-sensitive and drug-resistant ovarian cancer cells, transcript profiling revealed increased expression of RhoA in both CisR, PacR and dual cells, but near identical levels of RhoC (**Fig. 5**). Downstream of RhoA, ROCK II, associated with myosin II-dependent contractility (Yoneda et al. 2005), was upregulated. ROCK is known to activate myosin II by phosphorylating myosin light chain kinase (MLC) and inhibiting myosin phosphatase(Amano et al. 1996). The two non-muscle isoforms of non-muscle myosin II (NMM IIA and IIB) which have both been implicated in cancer invasion (Saha et al. 2011), were both upregulated in drug-resistant cells (**Fig. 5**).

**Figure 5:**
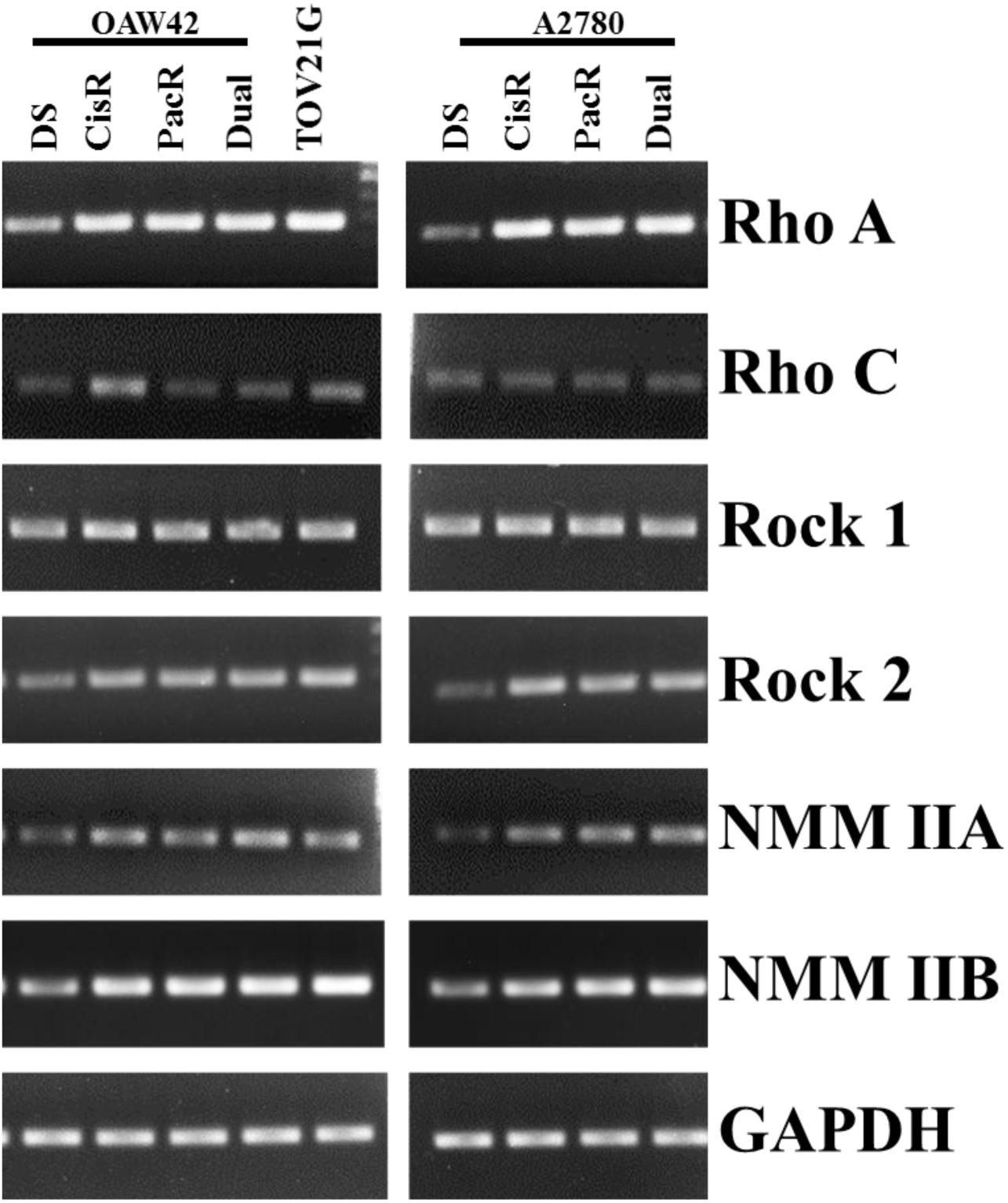
Molecular profiling of contractility-modulating proteins in drug-sensitive and drug-resistant ovarian cancer cells. Comparative analysis of mRNA levels of contractility proteins in drug-resistant and drug-sensitive cells assessed using RT-PCR. GAPDH proteinwas used as loading control.

In order to probe the localization of NMM IIA and NMM IIB proteins in drug-resistant ovarian cancer cells, we performed immunofluorescence staining. While NMM IIA showed peripheral localization, NMM IIB was seen to be present in the perinuclear region (**Fig. S4A**). Previous studies have suggested distinct roles of NMM IIA and NMM IIB, with NMM IIB mediating nuclear translocation through 3D matrices (Thomas et al. 2015).Given the perinuclear localization of NMM IIB observed in drug-resistant cells, we probed its role in modulating the distinct modes of invasion. We established stable knockdowns of NMM IIBinOAW42 dual drug-resistant cells (O-Dual) and in naturally cisplatin-resistant TOV21G cells. Experiments could not be performed with dual drug-resistantA2780 cells as NMM IIB knockdown led to cell death. shRNA based NMM IIB depletion was confirmed by western blot analysis with ~70-80% reduction in NMM IIB levels in both O-Dual and TOV21G cells (**Fig. 6A**). NMMIIA levels were unchanged in NMM IIB knockdown cells. Knockdown cells exhibited lower levels of pMLC (**Fig. 6A**) and vinculin (**Fig. S4B,C**)and were significantly softer compared to control cells (**Fig. 6C**).Further, sandwich invasion assays revealed inhibition of lateral invasion in knockdown cells (**Fig. 6D, Supp. Videos. 5-6**).

**Figure 6:**
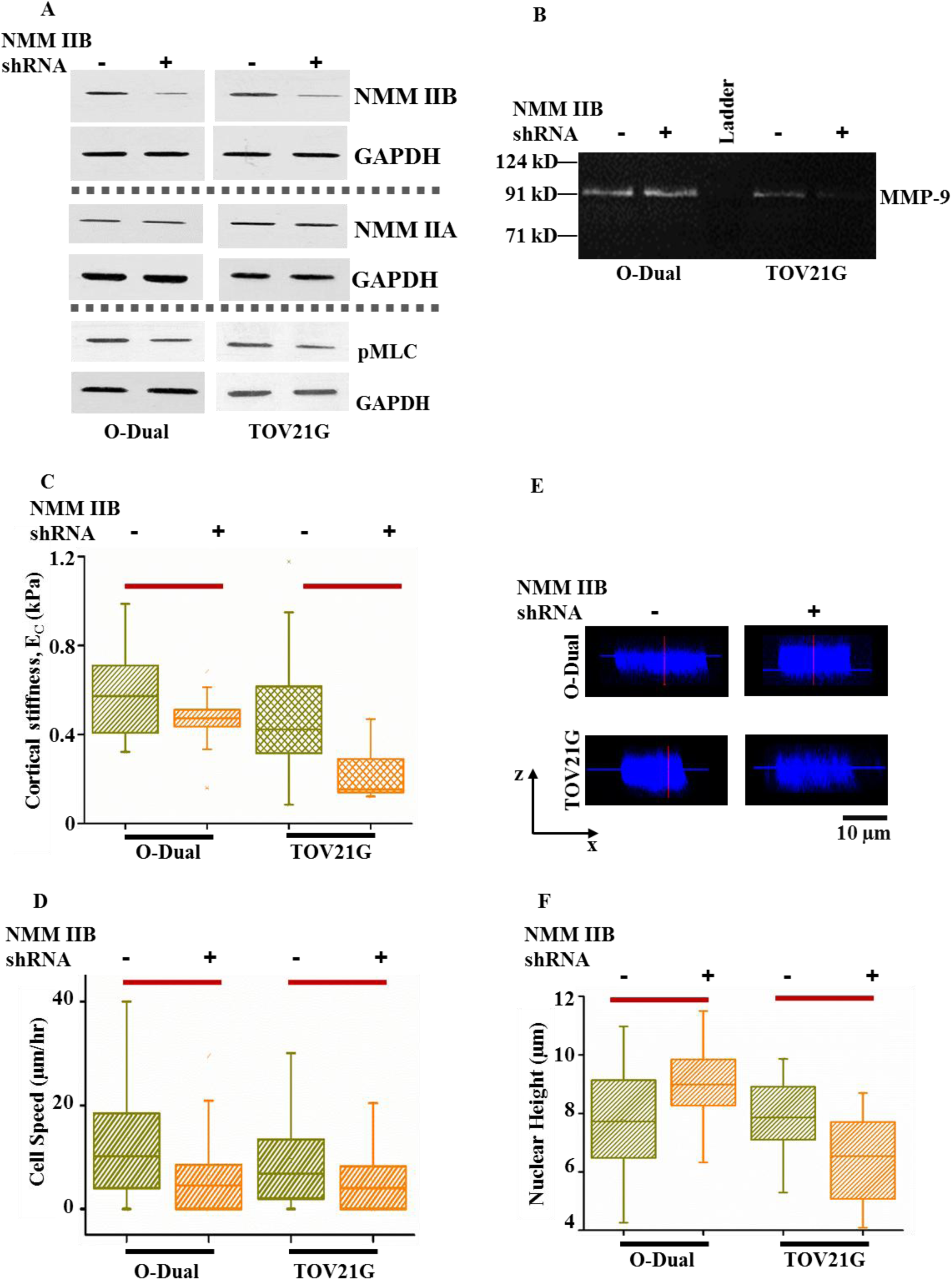
Non-muscle myosin IIB knockdown impairs cell migration through 3D matrices in drug-resistant ovarian cancer cells. **(A)** Validation of NMM IIB knockdown in dual-OAW42 and TOV21G cells. Western blots of NMM IIB, NMM IIAand pMLC in control and knockdown cells. GAPDH was used as a loading control. NMM IIB knockdown did not affect NMM IIA levels.**(B)**Influence of myosin IIB knockdown on MMP activity. Gelatin zymography was performed using conditioned media from control and knockdown cells. Loss of MMP-9 activity was seen in TOV21G knockdown cells but not in dual-OAW42 cells in which MMP-9 levels remained unperturbed.**(C)**Cortical cell stiffness of control and knockdown cells. Data represented as mean ± SEM (n = 100 cells for each condition, experiment repeated thrice). Red bars indicate statistical significance (*p < 0.05). **(D)** Quantification ofcell speed of control and knockdown cells in 1 mg/ml collagen sandwich gels. (n = 30cellsper condition across three independent experiments), Red bars indicate statistical significance (*p< 0.05) assessed using Student’s t test. **(E)** Representative x-z confocal images of DAPI stained nucleus in control and knockdown cells. **(F)** Quantification of nuclear height of control and knockdown cells (n = 30 cell nuclei across two independent experiments were analyzed for each condition). Error bar indicates SEM. Red bars indicate statistical significance(*p < 0.05).

To probe the mechanism by which NMM IIB regulates invasiveness of O-Dual and TOV21G cells, which utilize two distinct modes of invasion (i.e., protease independent versus protease dependent), zymography experiments were performed. While MMP-9 activity was unchanged in O-Dual cells upon NMM IIB knockdown, MMP-9 activity was significantly suppressed in TOV21G, suggesting NMM IIB regulates invasiveness of TOV21G by modulating MMP-9 activity (**Fig. 6B**). To next test if NMM IIB mediated invasiveness of O-Dual cells by effecting nuclear squeezing through pre-existing pores, nuclear height was compared in control and knockdown cells. Consistent with a nuclear squeezing role, nuclear height was increased in O-Dual knockdown cells compared to control (**Fig. 6E, F**). In contrast, nuclear height decreased in TOV21G knockdown cells. Together, our results demonstrate two distinct functions of NMM IIB in modulating cancer invasion either by regulating MMP-9 activity or by effecting nuclear squeezing (**Fig. 7**).

**Figure 7:**
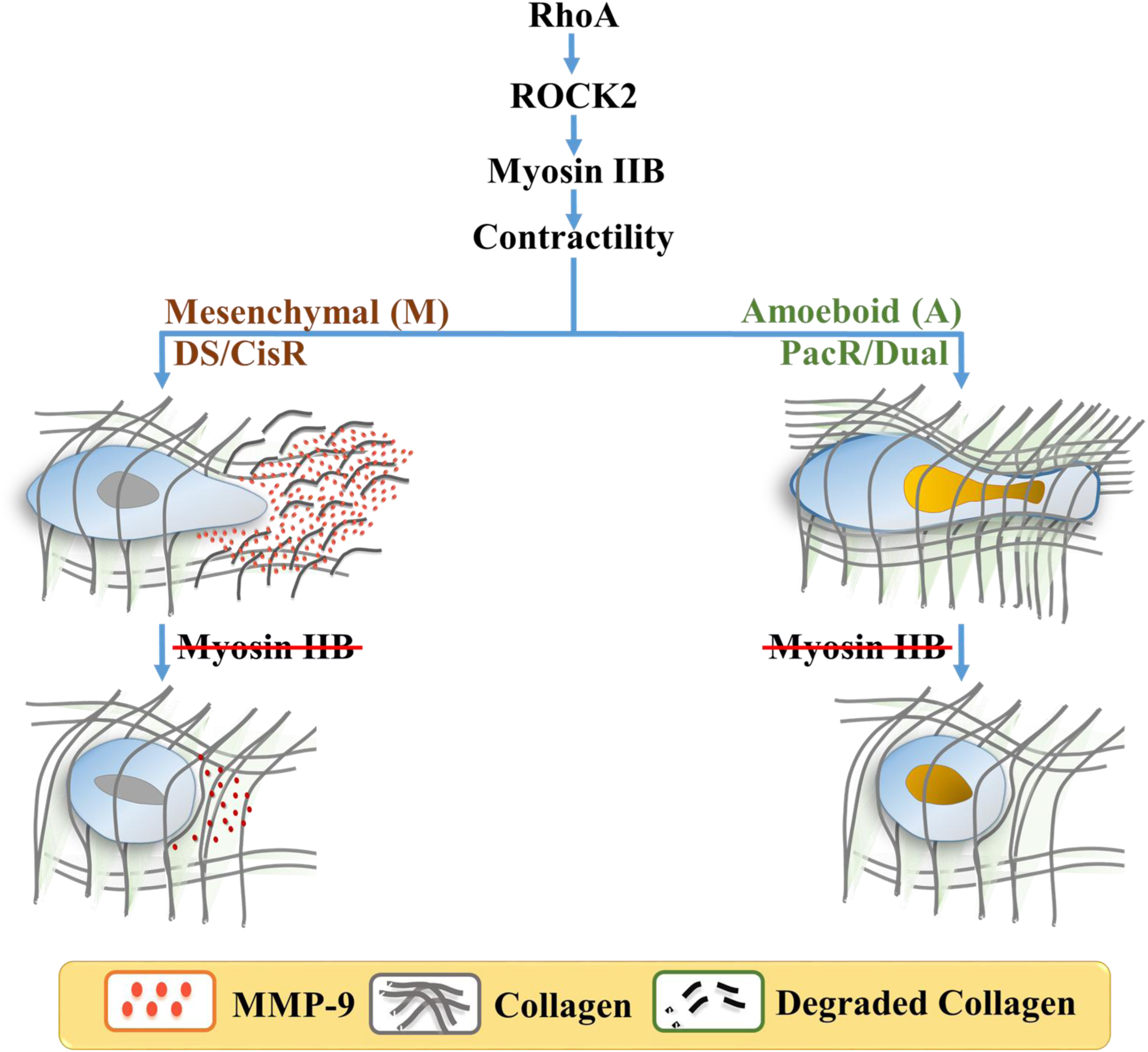
Model proposing regulation of invasion of drug-sensitive and drug-resistant ovarian cancer cells by myosin IIB. Cell contractility is controlled by the RhoA-ROCK2-myosin IIB signaling pathway. While DS and CisR cells utilize protease-dependent mesenchymal mode of invasion, PacR and dual cells utilize actomyosin contractility-dependent amoeboidal mode of invasion. Myosin IIB mediates MMP secretion in mesenchymal cells and nuclear squeezing in amoeboidal cells.

## Discussion

Different types of drug-resistant cancer cells such as prostate cancer and pancreatic cancer cells are known to become metastatically aggressive after undergoing EMT (epithelial to mesenchymal transition) (Thiery et al. 2009; Kajiyama et al. 2007; Kim et al. 2013). During mesenchymal migration, cells utilize proteases to degrade their surrounding matrix (Thiery et al. 2009). In MDA-MB-231 breast cancer cells and HT-1080 fibrosarcoma cells, inhibition of protease activity by GM6001 has been shown to cause a switch from protease-dependent to protease-independent mode of migration (Wolf et al. 2003). In chemotherapeutic drug-treated cells, drugs like paclitaxel may impart this effect by disrupting microtubular vesicular trafficking and inhibiting the release of MMP-2/9 as reported in melanoma cells (Schnaeker et al. 2004). This raises the possibility that drug-resistant cells may alter the mode of invasion depending on the molecular mechanism of operation of the drug. To address this question we studied the effect of acquisition of drug-resistance against two widely used chemotherapeutic drugs cisplatin and paclitaxel on epithelial ovarian cancer cell invasion.

We used serous-grade carcinoma cell lines (OAW42 and A2780) to develop cisplatin, paclitaxel and paclitaxel-cisplatin dual drug-resistant models and clear cell carcinoma cells (TOV21G) which are naturally cisplatin-resistant for our study. Serous carcinoma comprise of ~80% of detected ovarian cancers while clear cell carcinoma contribute to ~10-13% of ovarian cancers. Given the variation in morphology of drug-sensitive and drug-resistant cells, we anticipated differences in their invasion mechanism. Surprisingly, despite morphometric differences in cell size, we found similarity in their response to drug treatment. Cisplatin-resistance in both serous grade carcinoma (A2780 and OAW42) and naturally cisplatin-resistant clear cell carcinoma cells (TOV21G) caused cells to use protease-dependent mesenchymal mode of migration while paclitaxel-resistance or paclitaxel-cisplatin dual-resistance in serous grade carcinoma cells (A2780 and OAW42) disabled protease activity in them. Inhibition of protease activity in paclitaxel and paclitaxel-cisplatin dual-resistant cells lead to acquisition of contractility-dependent amoeboid mode of invasion. Based on comparative analysis of biophysical and molecular characteristics of drug-resistant ovarian cancer cells, for the first time, our data suggests that paclitaxel and paclitaxel-cisplatin-resistant ovarian cancer cells undergo MAT (mesenchymal to amoeboid transition).

Cortical softening represents a common feature across the different drug-resistant cells. Previously, a comparison of cortical stiffness of a range of different ovarian cancer cells have linked cell softness with increased metastatic potential (Xu et al. 2012). Another study of mechanical characterization of patient tumor cells and cancer cells using a magnetic tweezer system has also established a negative correlation between cell stiffness and invasion potential (Swaminathan et al. 2011). Our finding that paclitaxel and dual drug-resistant ovarian cancer cells are significantly softer than drug-sensitive ovarian cancer cells is consistent with these reports. Though we have found cisplatin-resistant cells also to be softer than drug-sensitive cells, recent studies with cisplatin-resistant ovarian cancer cells have reported the reverse trend of increased cell stiffness with drug resistance (Sharma et al. 2014). The divergence of our findings with this study may have arisen due to several factors. These differences may be attributed to differences in the type of AFM probe used, the location of the cell probed by the AFM and the indentation depth used for fitting the data (Meza et al. 2011; Radmacher 2007; Solon et al. 2007), the extent of cytoskeletal remodeling in cisplatin-resistant cellswhichisshown to depend on drug dosage (Seo et al. 2015).

In most adherent cells including fibroblasts and mesenchymal stem cells, cortical mechanics and contractile mechanics are closely coupled, i.e., increased contractility leads to increased cortical stiffness. However, this coupling is absent in embryonic stem cells(Poh et al. 2010). Here, we observe similar decoupling of contractile mechanics from cortical mechanics in drug-resistant cells, which are more contractile than drug-sensitive cells, but softer. Though NMM IIB knockdown cells were softer than control cells, they were inefficient in their invasiveness, suggesting that softening alone is not sufficient to promote invasiveness, and contractility is essential. Enhanced contractility is modulated by the Rho-ROCK pathway, specifically RhoA and ROCK2, which are known to regulate the assembly of central stress fibers(Totsukawa et al. 2004). Similar increase in contractility has been reported in paclitaxel-resistant prostate cancer cells(Kim et al. 2013).

Upregulation of both non-muscle isoforms of non-muscle myosin (NMM IIA and NMMIIB) was observed in drug-resistant cells. NMM IIA and NMM IIB have previously been shown to have non-redundant roles in cell migration. In MDCK epithelial kidney cells plated on 2D surfaces, NMM IIA is known to localize at the leading edge of the cells where they regulate focal adhesion dynamics(Jorrisch et al. 2013);NMM IIB on the other hand is found at the trailing end where they generate contractile forces for cell retraction(Jorrisch et al. 2013; Thomas et al. 2015). In line with existing literature, we noticed lamellar localization of NMM IIA and perinuclear localization of NMM IIB. During 3D cell migration, NMM IIB assists in nuclear squeezing, which is the rate limiting step in cell migration while moving through confined matrices (Thomas et al. 2015; Beadle et al. 2008). Moreover, enhanced metastatic potential of breast cancer stem cells has been correlated with elevated levels of NMM IIB, compared to non-cancer stem cells (Thomas et al. 2014).Motivated by these findings, we specifically probed the contributions of NMM IIB in drug-resistant cells, and found two distinct mechanisms by which NMM IIB regulates invasiveness. In paclitaxel and dual resistant cells, which migrate in protease independent manner, NMM IIB was found to mediate nuclear squeezing. However, in cisplatin-resistant cells, NMM IIB was found to regulate invasiveness by modulating protease activity. Though the mechanism has not been explored, it is possible that this regulation might be achieved by modulation of vesicular transport through actin-microtubule cooperation (Even-Ram et al. 2007). The reduction in nuclear height observed in NMM IIB knockdown cells may be due to increased coupling with the NMM IIA cytoskeleton which pulls the nucleus from the front(Petrie et al. 2014; McGregor et al. 2016).

In conclusion, our results demonstrate the existence of drug-type dependent distinct modes of invasion (mesenchymal versus amoeboidal) in drug-resistant ovarian cancer cells. Despite differences in their mechanism of invasion all drug-resistant ovarian cancer cells are regulated by NMM IIB-mediated actomyosin contractility. This raises the exciting possibility of suppressing cancer invasion by targeting NMM IIB.

